# Simultaneous GCaMP6-based Fiber Photometry and fMRI in Rats

**DOI:** 10.1101/146779

**Authors:** Zhifeng Liang, Yuncong Ma, Glenn D.R. Watson, Nanyin Zhang

**Affiliations:** Department of Biomedical Engineering, The Pennsylvania State University, University Park, PA 16802; The Huck Institutes of Life Sciences, The Pennsylvania State University, University Park, PA, 16802; Department of Psychology and Neuroscience, Duke University, Durham, NC 27708; Institute of Neuroscience, Chinese Academy of Sciences, Shanghai, China, 200031

**Keywords:** GCaMP6, fMRI, rats, fiber photometry

## Abstract

Understanding the relationship between neural and vascular signals is essential for interpretation of functional MRI (fMRI) results with respect to underlying neuronal activity. Simultaneously measuring neural activity using electrophysiology with fMRI has been highly valuable in elucidating the neural basis of the blood oxygenation-level dependent (BOLD) signal. However, this approach is also technically challenging due to the electromagnetic interference that is observed in electrophysiological recordings during MRI scanning. Recording optical correlates of neural activity, such as calcium signals, avoids this issue, and has opened a new avenue to simultaneously acquire neural and BOLD signals. The present study is the first to demonstrate the feasibility of simultaneously and repeatedly acquiring calcium and BOLD signals in animals using a genetically encoded calcium indicator, GCaMP6. This approach was validated with a visual stimulation experiment, during which robust increases of both calcium and BOLD signals in the superior colliculus were observed. In addition, repeated measurement in the same animal demonstrated reproducible calcium and BOLD responses to the same stimuli. Taken together, simultaneous GCaMP6-based fiber photometry and fMRI recording presents a novel, artifact-free approach to simultaneously measuring neural and fMRI signals. Furthermore, given the cell-type specificity of GCaMP6, this approach has the potential to mechanistically dissect the contributions of individual neuron populations with respect to BOLD signal, and ultimately reveal its underlying neural mechanisms.

## Introduction

Since its advent over two decades ago, functional MRI (fMRI) has been widely used as a non-invasive functional brain mapping technique in both human and animal studies. Typically, fMRI measures the blood oxygenation-level dependent (BOLD) signal, which is generally believed to be spatiotemporally linked to neural activity through the mechanism of neurovascular coupling (Hillman, 2014; Kim and Ogawa, 2012; Lecrux and Hamel, 2016). Neurovascular coupling is a highly complex physiological process that involves many types of brain cells, including excitatory pyramidal neurons, interneurons, astrocytes, pericytes, endothelia cells and smooth muscles. Sophisticated interplay among the activity of these cells triggers a wide range of downstream effects, which ultimately lead to changes in BOLD signal. In addition, this coupling between neural activity and vascular change likely varies in different brain regions (Thomsen et al., 2004), in different developmental stages (Kozberg et al., 2013) and pathological states (Iadecola, 2004).

Despite extensive research on this topic, the field is still far from a comprehensive and quantitative understanding of the neural basis of BOLD signal. This knowledge gap can be attributed to not only the complex nature of neurovascular coupling, but also to limitations in methods available to study this process. The commonly employed approach of simultaneous electrophysiology and fMRI recordings is technically challenging (Im et al., 2006; Liu et al., 2009), as it suffers from inherent electromagnetic interference from MRI scanning during electrical recording (Logothetis et al., 2001; Logothetis and Wandell, 2004). This interference also makes it difficult to differentiate neural activity from separate neuron populations, which limits the ability to elucidate the cellular mechanisms underlying neurovascular coupling.

The recent development of the optical fiber photometry technique provides a relatively simple and artifact-free method for simultaneously recording neural and BOLD signals (Schulz et al., 2012). Fiber photometry measures a single optical signal from a group of neurons, in a manner similar to an electrophysiological recording using an electrode. However, owing to its simplicity and MR compatibility, it can be implemented with the tight space limitation and non-ferromagnetic requirement inside MRI scanners. A previous study utilized chemical dyes (OGB-1, Rhod-2 and Fluo-4) as calcium indicators to simultaneously acquire calcium and BOLD signals in rats (Schulz et al., 2012), which showed robust activation of both signals in a forepaw stimulation paradigm. Another recent study employed a similar approach, with the neuronal activity modulated using optogenetics (Schmid et al., 2016). However, due to the nature of acute loading and decay of chemical dyes in brain tissue, it is difficult to conduct repeated and longitudinal experiments with chemical dyes. Moreover, cell-type specificity of the calcium signal is limited by the chemical properties of dyes.

To overcome these limitations, we developed a setup that allows simultaneous GCaMP6-based fiber photometry and fMRI data acquisition in rats. In this setup, GCaMP6 (Chen et al., 2013), a newly improved genetically encoded calcium indicator (GECI), was used to replace chemical dyes. As the calcium sensitivity for GCaMP6 lasts for months, this change allows for repeated measurement in the same animal. In addition, well-established genetic tools allow GCaMP6 to be expressed in a specific neuron type (e.g. excitatory or inhibitory neurons). This cell-type specificity makes it possible to further dissect the contributions of individual neuron populations with respect to fMRI signal in the future. With a visual stimulation paradigm, the present study is the first to demonstrate the feasibility and reliability of simultaneous GCaMP6-based fiber photometry and fMRI data acquisition in rats.

## Methods

### Animals

Seven adult male Long-Evans rats (300-500g, Charles River Laboratories, Wilmington, MA) were housed in Plexiglas cages and maintained on a 12 h light:12 h dark schedule in a room temperature of 22-24° C. Food and water were provided ad libitum. All procedures were approved by the Institutional Animal Care and Use Committee of the Pennsylvania State University.

### Surgery and Extracellular neuronal recordings

Rats were anesthetized by intramuscular (IM) injections of ketamine (40 mg/kg) and xylazine (12 mg/kg). Dexamethasone (0.5 mg/kg) was injected IM to prevent tissue inflammation. All rats were intubated, placed in a stereotaxic frame (David Kopf Instruments, Tujunga, CA), and artificially ventilated with oxygen. During surgery, the animal’s body temperature was maintained at 37° C and its heart rate and SpO_2_ were monitored. A small craniotomy was made unilaterally over the right superior colliculus (6.3 mm rostral and 0.8 mm lateral to bregma).

Prior to AAV injection, recordings in the medial superior colliculus were made with a hypertonic saline filled glass pipette to determine the depth of visually responsive neurons. Visual stimulation was administered by a blue laser source (100 mW, 473 nm, Opto Engine) through an optic fiber positioned ~5 cm away from the contralateral eye. On each computer-controlled trial, a single 25 ms blue light flash was administered. Neuronal discharges were recorded as previously described (Liang et al., 2015). Discharges with a signal-to-noise ratio of a least 3:1 were time-stamped at a resolution of 0.1 ms and were displayed as peristimulus-timed histograms (PSTHs). After measuring the mean rate of spontaneous activity recorded before stimulus onset, the 99% confidence limits were calculated for each PSTH, and visually-evoked responses were considered significant if they exceeded the confidence limits.

After robust visual responses were recorded (typically between 2.6 to 2.8 mm deep), AAV5.Syn.GCaMP6s (800-1000nl, Penn Vector Core) was injected at 50 μm and 250 μm below the electrophysiology recording depth through a micropipette syringe fitted with a glass pipette tip (Hamilton Company, Reno, NV). After the syringe was withdrawn, an optic fiber (0.4 mm diameter) embedded in a 2.5 mm ceramic ferrule (Thorlabs, Newton, NJ) was slowly advanced towards the injection site until it reached 250 μm above the electrophysiology recording depth. Three plastic screws (0.08 × 0.125, nylon; PlasticsOne, Roanoke, VA) or ceramic screws (0.08 × 0.125; Ortech, Sacramento, CA) were placed along the temporal ridge. Dental cement was applied around the screws and ceramic ferrule to form a platform. The rat was placed back into its home cage for at least 4 weeks to allow for recovery and GCaMP expression.

### Simultaneous calcium-based fiber photometry and fMRI

Rats were anesthetized using dexmedetomidine (initial bolus injection of 0.05 mg/kg followed by subcutaneous infusion at 0.1 mg/kg/h 10 minutes after the bolus). The fiber photometry setup was adopted from Gunaydin et al. (Gunaydin et al., 2014) with minor modifications. Briefly, GCaMP was excited by a 473nm laser (Opto Engine LLC, Midvale, UT) chopped at 400 Hz with an optical chopper (Thorlabs, Newton, NJ), reflected off a dichroic mirror (FF495, Semrock, Rochester, New York), then coupled into an optical patch cable (0.4mm diameter, 7m long) using a 40×0.65 NA microscope objective (Olympus) and a fiber launch (Thorlabs, Newton, NJ). The laser power at the tip of implanted optical fibers was approximately 2mW. The emitted fluorescent GCaMP signal was collected through the same optical cable, objective and dichroic mirror, then filtered by a bandpass filter (FF01-520/35, Semrock, Rochester, New York), and finally collected by a silicon photomultiplier (MiniSM 30035, SensL, Ireland). The output of the silicon photomultiplier was amplified by a lock-in amplifier (SR810, Stanford Research Systems, Sunnyvale, CA) with a time constant of 3 ms, and then digitized using an NI DAQ device (NI USB-6211, National Instruments, Austin, TX) at 100 Hz with custom-written LabView programs. The acquisition of the optical signal was triggered by a TTL signal from the MRI scanner at the beginning of each fMRI run, which ensured that the optical signal was synchronized with fMRI data. The optical setup was housed in a separate room adjacent to the MRI scanner.

MRI experiments were conducted on a 7T system with a Bruker console (Billerica, MA). During each MRI session, anatomical images were first acquired using a T1 weighted RARE sequence (matrix size = 256×256; FOV = 3.2×3.2 cm; slice number = 15; slice thickness = 1 mm). Gradient-echo fMRI images were acquired using echo-planar imaging (EPI) with the following parameters: repetition time (TR) = 1 s; echo time (TE) = 15 ms; matrix size = 64×64; FOV = 3.2×3.2 cm; slice number = 15; and slice thickness = 1 mm. The setup of simultaneous calcium-based fiber photometry and fMRI is schematically illustrated in Figure 1. An exemplar EPI volume is shown in Supplemental Figure 1.

**Figure 1.**
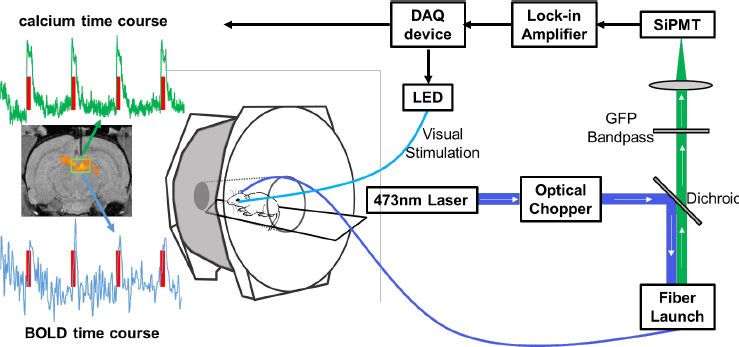
Schematic illustration of the setup of simultaneous calcium-based fiber photometry and fMRI. Center: Anesthetized rodent in bore of MRI tethered to optic patch cable. Right: Fiber photometry set-up. GCaMP is excited by a 473 nm laser chopped with an optical chopper reflected off a dichroic mirror (blue line). This signal is coupled into an optical patch cable using a microscope objective and a fiber launch. The emitted fluorescent GCaMP signal is then collected through the same optical apparatus, filtered by a GFP bandpass filter, and collected by a silicon photomultiplier (SiPMT) (green line). Top: The output of the SiPMT is amplified by a lock-in amplifier and digitized using a DAQ device. Visual stimulation is controlled by the same DAQ device to drive an LED light source. Left: Acquired data from simultaneous calcium-based fiber photometry and fMRI. Both calcium and BOLD time courses are obtained from a region of interest under the optic fiber. T1 weighted fMRI image shows implanted optic fiber with respect to a group functional activation map. Red line denotes visual stimulation of the contralateral eye.

Visual stimulation (5Hz, 2s, 5s or 10s with inter-stimulus intervals of 15s, 25s or 50s, respectively) was controlled by the same NI DAQ device used for optical signal collection. An optical fiber was coupled with a blue LED and the tip of the optical fiber was placed in front of the rat’s left eye. Each simultaneous fiber photometry and fMRI session consisted of 2-3 scans, and each scan consisted of 10, 7 or 5 epochs for 2s, 5s or 10s of stimulation, respectively.

### Histology

Each rat was deeply anesthetized with intramuscular injections of ketamine (80 mg/kg) and xylazine (6 mg/kg), and then was perfused transcardially with heparinized saline, 4% paraformaldehyde, and 4% paraformaldehyde in 10% sucrose. The brain was removed and stored in 4% paraformaldehyde with 30% sucrose in a refrigerator for 3-5 d. A freezing microtome was used to cut each brain into serially ordered 60 μm thick sections that were placed in 0.1 M PBS. A 1-in-2 series of sections were processed and mounted onto gel-dipped slides in serial order. The first series, used to visualize cytoarchitecture, was processed for the presence of cytochrome oxidase. A second series was used to visualize fluorescence to confirm AAV expression and the injection site, and did not undergo any histochemical processing. Mounted sections in each series were dehydrated in ethanol, defatted in xylene, and coverslipped with cytoseal.

### Data analysis

The calcium signal was low pass filtered (cutoff: 10 Hz), detrended and then converted to a percentage change. For imaging data, anatomical images were first manually aligned to a fully segmented rat brain atlas embedded in Medical Image Visualization and Analysis (MIVA, ccni.wpi.edu). EPI images were motion-corrected, spatially smoothed (FWHM 1mm), and then subjected to statistical analysis using SPM12 (http://www.fil.ion.ucl.ac.uk/spm/). The significance level was set at p < 0.05, after false discovery rate (FDR) correction (Genovese et al., 2002). The first five volumes of each EPI scan were discarded prior to data analysis to allow magnetization to reach steady state.

To plot BOLD and calcium signals together, both signals were re-sampled to 10 Hz. For each stimulation paradigm, the BOLD time course was obtained by averaging the time courses of significantly activated voxels around the tip of the optical fiber in three consecutive slices (i.e. the slice with the implanted optical fiber, as well as one anterior slice and one posterior slice) for each rat. All data analyses were conducted with custom-written Matlab (MathWorks, Natick, MA) scripts unless otherwise noted.

## Results

The experimental design is summarized in Figure 2. During surgery, all animals showed a robust electrophysiological response to light flashes determined by extracellular recordings (Figure 3A). In addition, the expression of GCaMP was verified histologically after all functional experiments were concluded (Figure 3C).

**Figure 2.**
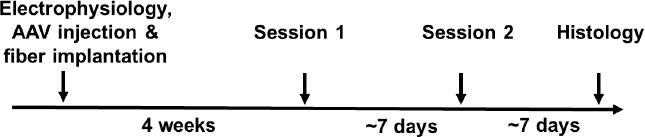
Summary of experimental design.

**Figure 3.**
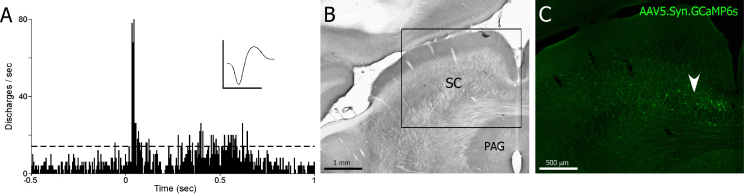
Injection of AAV5.Syn.GCaMP6s into visually responsive superior colliculus. **A,** PSTH shows neural responses in medial superior colliculus during a 25ms light flash. Dashed line indicates 99% confidence limit. Mean waveform scale: 1ms, 500 mV. PSTH: 100 trials, 10 ms bin widths. **B,** Cytochrome oxidase processed coronal section through the superior colliculus. Inset shown in C. **C,** Fluorescent image of transfected neurons in the deep layers of the medial superior colliculus after injection of AAV5.Syn.GCaMP6s (Penn Vector Core). White arrow denotes injection area where visually responsive neurons were recorded in A.

During the simultaneous fiber photometry and fMRI experiment, we observed robust increases of calcium and BOLD signals in response to visual stimulation (Figure 4). Light flashes induced a significant increase of BOLD signal in the superior colliculus (Figure 4A). Importantly, the simultaneously acquired calcium signal from the same brain area also displayed clear activation that was temporally synchronized with the corresponding stimulation paradigm. Figure 4B displays the averaged time courses of the BOLD and calcium signals around the tip of the optical fiber (5 rats, 2-sec stimulation). As calcium signal measures spiking activity, the onset and peak times of GCaMP signal were appreciably faster than the BOLD response, due to the hemodynamic delay.

**Figure 4.**
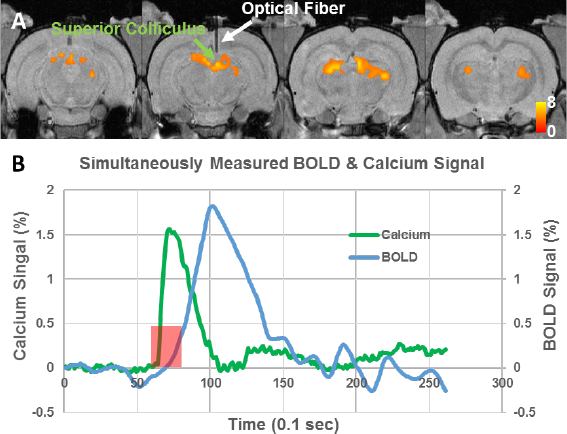
**A**, Group functional activation map during visual stimulation (N=7). T values were thresholded at p<0.05 (FDR corrected). White arrow indicates implanted optical fiber. **B,** Averaged time courses of the BOLD and calcium signals around the tip of the optical fiber (5 rats, 2-sec stimulation).

We found that both BOLD and calcium signals were reliably detected in all animals regardless of the stimulation duration. Figure 5A demonstrates reliable BOLD and calcium responses during each individual trial in one animal. Figures 5B-H display averaged BOLD and calcium time courses for each animal. The calcium signal from all animals exhibited an onset within several hundred milliseconds, a slight adaptation during stimulation-on periods, which was more prominent in 5-s or 10-s stimulation paradigms, and a fast decrease after the stimulation offset. Interestingly, the peak amplitude of the BOLD signal showed less inter-subject variability relative to the calcium signal, which could vary from 1% to 10% across animals.

**Figure 5.**
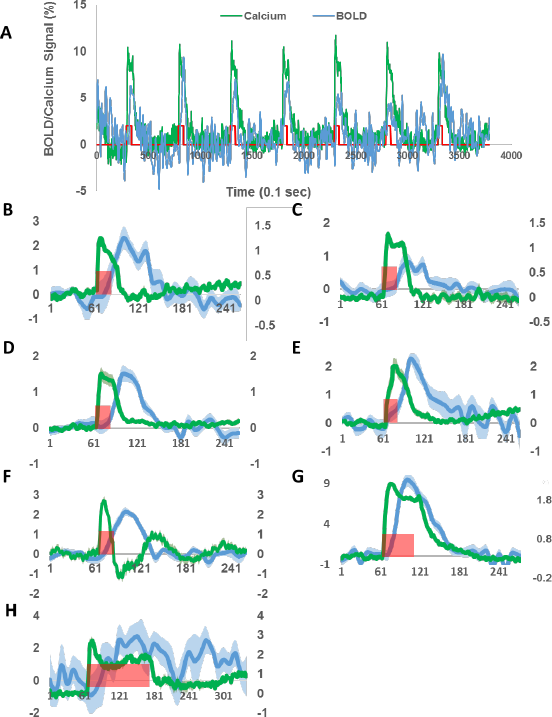
Simultaneously acquired BOLD and calcium time courses in each individual rat. **A,** BOLD and calcium responses to single trials of visual stimulation. **B-H,** Averaged BOLD and calcium time courses for Rat 1 to 7, respectively. Green traces: calcium signals. Blue traces: BOLD signals. Shaded regions denote standard error of the mean. Red bars indicate visual stimulation periods. X axis: time in 0.1s. Left y axis: % change of BOLD signal. Right y axis: % change of calcium signal.

As the GCaMP signal was picked up by a chronically implanted optical fiber, this setup makes it possible to repeatedly and longitudinally measure calcium and BOLD signals in the same animal. To demonstrate this advantage, Figure 6 shows BOLD and calcium signals in two rats that were imaged twice, separated by approximately one week. Reproducible BOLD and calcium activations were observed in both animals, with similar temporal patterns between the two sessions. Interestingly, the calcium peak amplitude of the second sessions was higher compared to the first sessions, possibly due to continuous GCaMP expression over time.

**Figure 6.**
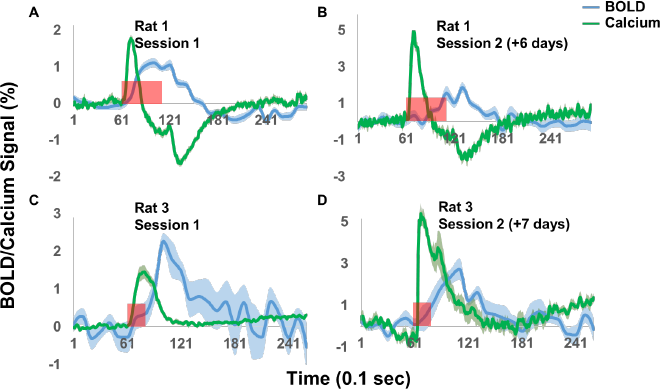
Repeated simultaneous measurement of BOLD and calcium signals. Green traces: calcium signals. Blue traces: BOLD signals. Shaded regions denote standard error of the mean. Red bars indicate visual stimulation periods. X axis: time in 0.1s. Y axis: % signal change.

## Discussion

To the best of our knowledge, this is the first study to demonstrate the feasibility of simultaneously measuring fMRI and calcium signals using GCaMP6-based fiber photometry. When comparing to previously used fluorescent chemical dyes (Schmid et al., 2016; Schulz et al., 2012), the utilization of GCaMP6 (Chen et al., 2013), a newly improved protein-based GECI, enabled repeated measurement of calcium and BOLD signals. In addition, as a GECI, GCaMP6 can be selectively expressed in a defined cell population, which allows neural activity with cell-type specificity to be measured. This ability makes it possible to dissect the contributions of activities from individual neuron populations to the fMRI signal, which will help ultimately decipher its neural basis.

It is generally assumed that BOLD signal is spatially and temporally linked to local neuronal activity (Liu et al., 2010; Zhang et al., 2008a; Zhang et al., 2009; Zhang et al., 2008b), and such neurovascular coupling serves as the fundamental basis of BOLD fMRI (Liu et al., 2010; Roy and Sherrington, 1890; Zhang et al., 2008a; Zhang et al., 2009; Zhang et al., 2008b). Due to the prominent role of BOLD fMRI in functional brain mapping, a large body of research is aimed at unraveling the mechanisms of neurovascular coupling (Hillman, 2014; Kim and Ogawa, 2012). In this research field, a commonly employed technique is simultaneous electrophysiology-fMRI acquisition. For example, by utilizing simultaneous extracellular electrical recording and fMRI, the seminal work of Logothetis et al. (Logothetis et al., 2001) showed that local field potentials were closely related to BOLD signal. Although feasible (Im et al., 2007; Im et al., 2006; Liu et al., 2009), simultaneous electrophysiology-fMRI acquisition is technically challenging as electrophysiology data suffer severe electromagnetic interference from MRI scanning, which significantly limits its applicability. In addition, this extracellular recording reflects the overall activity of multiple cell types, and it is not possible to separate out cell-type specific contributions to BOLD signal, which limits its ability to uncover the cellular-level mechanism of neurovascular coupling.

To avoid these issues, one emerging alternative approach is to record neural activity during fMRI scanning using an optical signal, which has virtually no MRI-related interference. Techniques of chemical dyes or genetically encoded optical indicators have been greatly expanded and improved in recent years, and have been extensively used to record calcium or voltage-sensitive signals (Weber and Helmchen, 2014). Notably, simultaneous fiber photometry and fMRI has been successfully demonstrated in rats (Schmid et al., 2016; Schulz et al., 2012). Fiber photometry records an optical signal from a neuronal population through an optical fiber chronically implanted in the animal. Such a simple setup makes it particularly suitable for use inside MRI scanners, as it requires minimal space and does not involve any metal components. By utilizing chemical dyes (OGB-1, Rhod-2 and Fluo-4) as calcium indicators, two studies simultaneously recorded the calcium signal, which directly reflects spiking activity, and fMRI data in animals (Schmid et al., 2016; Schulz et al., 2012). This novel method has opened a new avenue to investigating the neurovascular coupling relationship. Nonetheless, chemical dyes applied in the previous two studies have two major limitations. First, acute loading and decay of chemical dyes make longitudinal measurement very difficult. Second, although some dyes do have some tissue specificity (e.g., Rhod-2 and Fluo-4 can be specific to astrocytes), this specificity in general is rather limited and cannot be conveniently extended to other cell types.

These two limitations can be overcome by GECIs such as GCaMP6. To examine this notion, our study replaced chemical dyes with an Adeno-associated virus (AAV) that expresses GCaMP6, and used a chronically implanted optical fiber to collect the calcium signal. With this setup, we demonstrated the feasibility and validity of simultaneous GCaMP6-based fiber photometry and fMRI using a visual stimulation paradigm. First, we identified the area of the superior colliculus that responded to visual stimulation using electrophysiology recording outside of the MRI scanner. From the same area, both calcium and BOLD responses to light flashes were reliably detected (Figures. 4 and 5). BOLD fMRI results were consistent with previous rodent fMRI studies using a similar paradigm (Chan et al., 2010; Lau et al., 2011), and GCaMP expression was confirmed histologically. Notably, one out of seven rats showed a slightly different temporal pattern of calcium signal (Figs. 5F and 6A and B), with an undershoot after the initial peak. One possible explanation for this temporal pattern is that the location of the fiber tip in this rat was immediately adjacent to blood vessels, and the increase of cerebral blood volume (CBV) and oxy-hemoglobin led to more absorption of light at 473nm (i.e. the wavelength for GCaMP excitation). The exact mechanism underlying the post-stimulation undershoot of calcium signal in this particular rat needs further investigation. However, this outlier should not affect the validity of the technique reported in the current study. Taken together, our results indicate that combining GCaMP6-based fiber photometry and fMRI can reliably measure neural and fMRI signals simultaneously.

Importantly, the setup adopted in the present study allows convenient, longitudinal measurement of calcium and BOLD signals potentially for months, without the need of any additional procedures after the initial surgery. Our data show that both calcium and BOLD signals were reproducible for two measurements separated by a week. This advantage is time- and cost-efficient, and is also critical for paradigms that require longitudinal monitoring of neural and BOLD activities.

Another advantage offered by simultaneous GCaMP6-based fiber photometry and fMRI is cell-type specific recording of neural activity. Well-established genetic technology allows the expression of GCaMP6 to be controlled by viral vectors to achieve cell-type specificity. In the current study, GCaMP6 was ubiquitously expressed in all neurons under the control of the synapsin promoter, as our purpose is mainly technical validation. In future studies, GCaMP specificity for individual cell types can be conveniently achieved by cell-type specific promoters in viral vectors, or the widely used *Cre-loxP* strategy when combined with transgenic *Cre* knockin animals (Huang et al., 2014; Tsien, 2016). Therefore, the current approach, with unprecedented cell-type specificity, may help unravel contributions of individual cell populations to BOLD signal.

The approach established in the present study has also opened up opportunities in several other research areas. For example, a natural extension of the present study would be fiber photometry in multiple sites (Kim et al., 2016) in combination with fMRI, which will allow the examination of neurovascular coupling at the neural circuit level. With further improved optical fiber-based endoscopy methods (Murray and Levene, 2012), it can potentially bring the cellular resolution of optical imaging into MRI scanners. Other possibilities include replacing current GECIs with genetically encoded voltage indicators (GEVIs), which will make the capacity of fiber photometry comparable to electrophysiology (Yang and St-Pierre, 2016). Additionally, because of the chronic nature of the current approach, it can be easily extended to awake animals, as the setup in the animal was the same as our previous study of simultaneous optogenetics and fMRI in awake rats (Liang et al., 2015).

There are a few limitations in the current study. First, a negative control (e.g. viral vector expressing GFP only) should further strengthen the validity of the current results. Nonetheless, it is unlikely that non-calcium related signal of GCaMP6s could significantly affect the results, as the calcium signal changes upon visual stimulation were robust and the temporal pattern was in excellent agreement with the known GCaMP6s kinetics (Chen et al., 2013). Indeed, the fluorescent signal detected by the optical fiber during the stimulation-off time also served as controls. Second, in the current setup, the MR signal in functional images immediately adjacent to the implanted optical fiber was attenuated due to the magnetic susceptibility effect (Supplemental Figure 1). This technical challenge could be potentially addressed using other functional imaging sequences less susceptible to this effect, or using optical materials with magnetic property similar to the brain tissue. Third, it would be helpful to validate the calcium signal by simultaneously acquiring electrophysiology data. However, we believe that the current results are sufficient to establish the validity of calcium signals for two reasons: 1) Electrophysiology recording in response to visual stimulation was performed (see Figure 2) for every rat, and viral vector expressing GCaMP6s was injected at the same location, immediately followed by optical fiber implantation. This step ensured that the calcium signal detected was from the same location where the electrophysiology signal was recorded. 2) Calcium signal in response to visual stimulation is temporally synchronized with typical GCaMP6s kinetics. Additionally, the calcium signal from GCaMP is well characterized to be highly correlated with spiking activity. Therefore, we believe the current design and data are sufficient to establish the validity of calcium signal we observed.

## Summary

Overall, the current study established the method for simultaneous GCaMP6-based fiber photometry and fMRI in rats. This approach presents significant benefits over existing methods, including the ability of longitudinal measurement and potential cell-type specificity. By taking advantage of rapidly improving genetic manipulation techniques and genetically encoded optical indicators, this approach will be able to provide numerous possibilities for further improvement. As fMRI remains one of the most popular functional neuroimaging tools, the need of understanding its neural mechanism is ever pressing. The approach presented in the current study provides a novel tool for further dissecting neurovascular coupling.

## Acknowledgement

The work was supported by the National Institutes of Health (R01MH098003 and R01NS085200) awarded to Nanyin Zhang, with additional support from Institute of Neuroscience, Chinese Academy of Sciences to Zhifeng Liang.

## References

Chan, K.C., Xing, K.K., Cheung, M.M., Zhou, I.Y., Wu, E.X., 2010. Functional MRI of postnatal visual development in normal and hypoxic-ischemic-injured superior colliculi. NeuroImage 49, 2013–2020.

Chen, T.W., Wardill, T.J., Sun, Y., Pulver, S.R., Renninger, S.L., Baohan, A., Schreiter, E.R., Kerr, R.A., Orger, M.B., Jayaraman, V., Looger, L.L., Svoboda, K., Kim, D.S., 2013. Ultrasensitive fluorescent proteins for imaging neuronal activity. Nature 499, 295–300.

Genovese, C.R., Lazar, N.A., Nichols, T., 2002. Thresholding of statistical maps in functional neuroimaging using the false discovery rate. Neuroimage 15, 870–878.

Gunaydin, L.A., Grosenick, L., Finkelstein, J.C., Kauvar, I.V., Fenno, L.E., Adhikari, A., Lammel, S., Mirzabekov, J.J., Airan, R.D., Zalocusky, K.A., Tye, K.M., Anikeeva, P., Malenka, R.C., Deisseroth, K., 2014. Natural neural projection dynamics underlying social behavior. Cell 157, 1535–1551.

Hillman, E.M., 2014. Coupling mechanism and significance of the BOLD signal: a status report. Annu Rev Neurosci 37, 161–181.

Huang, Z.J., Taniguchi, H., He, M., Kuhlman, S., 2014. Cre-dependent adeno-associated virus preparation and delivery for labeling neurons in the mouse brain. Cold Spring Harb Protoc 2014, 190–194.

Iadecola, C., 2004. Neurovascular regulation in the normal brain and in Alzheimer’s disease. Nat Rev Neurosci 5, 347–360.

Im, C.H., Gururajan, A., Zhang, N., Chen, W., He, B., 2007. Spatial resolution of EEG cortical source imaging revealed by localization of retinotopic organization in human primary visual cortex. J Neurosci Methods 161, 142–154.

Im, C.H., Liu, Z., Zhang, N., Chen, W., He, B., 2006. Functional cortical source imaging from simultaneously recorded ERP and fMRI. J Neurosci Methods 157, 118–123.

Kim, C.K., Yang, S.J., Pichamoorthy, N., Young, N.P., Kauvar, I., Jennings, J.H., Lerner, T.N., Berndt, A., Lee, S.Y., Ramakrishnan, C., Davidson, T.J., Inoue, M., Bito, H., Deisseroth, K., 2016. Simultaneous fast measurement of circuit dynamics at multiple sites across the mammalian brain. Nat Methods 13, 325–328.

Kim, S.G., Ogawa, S., 2012. Biophysical and physiological origins of blood oxygenation level-dependent fMRI signals. J Cereb Blood Flow Metab 32, 1188–1206.

Kozberg, M.G., Chen, B.R., DeLeo, S.E., Bouchard, M.B., Hillman, E.M., 2013. Resolving the transition from negative to positive blood oxygen level-dependent responses in the developing brain. Proc Natl Acad Sci USA 110, 4380–4385.

Lau, C., Zhou, I.Y., Cheung, M.M., Chan, K.C., Wu, E.X., 2011. BOLD temporal dynamics of rat superior colliculus and lateral geniculate nucleus following short duration visual stimulation. PLoS One 6, e18914.

Lecrux, C., Hamel, E., 2016. Neuronal networks and mediators of cortical neurovascular coupling responses in normal and altered brain states. Philos Trans R Soc Lond B Biol Sci 371.

Liang, Z., Watson, G.D., Alloway, K.D., Lee, G., Neuberger, T., Zhang, N., 2015. Mapping the functional network of medial prefrontal cortex by combining optogenetics and fMRI in awake rats. Neuroimage 117, 114–123.

Liu, Z., Rios, C., Zhang, N., Yang, L., Chen, W., He, B., 2010. Linear and nonlinear relationships between visual stimuli, EEG and BOLD fMRI signals. Neuroimage 50, 1054–1066.

Liu, Z., Zhang, N., Chen, W., He, B., 2009. Mapping the bilateral visual integration by EEG and fMRI. Neuroimage 46, 989–997.

Logothetis, N.K., Pauls, J., Augath, M., Trinath, T., Oeltermann, A., 2001. Neurophysiological investigation of the basis of the fMRI signal. Nature 412, 150–157.

Logothetis, N.K., Wandell, B.A., 2004. Interpreting the BOLD signal. Annu Rev Physiol 66, 735–769.

Murray, T.A., Levene, M.J., 2012. Singlet gradient index lens for deep in vivo multiphoton microscopy. J Biomed Opt 17, 021106.

Roy, C.S., Sherrington, C.S., 1890. On the Regulation of the Blood-supply of the Brain. J Physiol 11, 85–158 117.

Schmid, F., Wachsmuth, L., Schwalm, M., Prouvot, P.H., Jubal, E.R., Fois, C., Pramanik, G., Zimmer, C., Faber, C., Stroh, A., 2016. Assessing sensory versus optogenetic network activation by combining (o)fMRI with optical Ca2+ recordings. J Cereb Blood Flow Metab 36, 1885–1900.

Schulz, K., Sydekum, E., Krueppel, R., Engelbrecht, C.J., Schlegel, F., Schroter, A., Rudin, M., Helmchen, F., 2012. Simultaneous BOLD fMRI and fiber-optic calcium recording in rat neocortex. Nat Methods 9, 597–602.

Thomsen, K., Offenhauser, N., Lauritzen, M., 2004. Principal neuron spiking: neither necessary nor sufficient for cerebral blood flow in rat cerebellum. J Physiol 560, 181–189.

Tsien, J.Z., 2016. Cre-Lox Neurogenetics: 20 Years of Versatile Applications in Brain Research and Counting. Front Genet 7, 19.

Weber, B., Helmchen, F., 2014. Optical imaging of neocortical dynamics. Humana Press; Springer, New York.

Yang, H.H., St-Pierre, F., 2016. Genetically Encoded Voltage Indicators: Opportunities and Challenges. J Neurosci 36, 9977–9989.

Zhang, N., Liu, Z., He, B., Chen, W., 2008a. Noninvasive study of neurovascular coupling during graded neuronal suppression. J Cereb Blood Flow Metab 28, 280–290.

Zhang, N., Yacoub, E., Zhu, X.H., Ugurbil, K., Chen, W., 2009. Linearity of blood-oxygenation-level dependent signal at microvasculature. Neuroimage 48, 313–318.

Zhang, N., Zhu, X.H., Chen, W., 2008b. Investigating the source of BOLD nonlinearity in human visual cortex in response to paired visual stimuli. Neuroimage 43, 204–212.

